# *mity*: A highly sensitive mitochondrial variant analysis pipeline for whole genome sequencing data

**DOI:** 10.1101/852210

**Authors:** Clare Puttick, Kishore R Kumar, Ryan L Davis, Mark Pinese, David M Thomas, Marcel E Dinger, Carolyn M Sue, Mark J Cowley

## Abstract

**Motivation:** Mitochondrial diseases (MDs) are the most common group of inherited metabolic disorders and are often challenging to diagnose due to extensive genotype-phenotype heterogeneity. MDs are caused by mutations in the nuclear or mitochondrial genome, where pathogenic mitochondrial variants are usually heteroplasmic and typically at much lower allelic fraction in the blood than affected tissues. Both genomes can now be readily analysed using unbiased whole genome sequencing (WGS), but most nuclear variant detection methods fail to detect low heteroplasmy variants in the mitochondrial genome.

**Results:** We present *mity*, a bioinformatics pipeline for detecting and interpreting heteroplasmic SNVs and INDELs in the mitochondrial genome using WGS data. In 2,980 healthy controls, we observed on average 3,166× coverage in the mitochondrial genome using WGS from blood. *mity* utilises this high depth to detect pathogenic mitochondrial variants, even at low heteroplasmy. *mity* enables easy interpretation of mitochondrial variants and can be incorporated into existing diagnostic WGS pipelines. This could simplify the diagnostic pathway, avoid invasive tissue biopsies and increase the diagnostic rate for MDs and other conditions caused by impaired mitochondrial function.

**Availability:** *mity* is available from https://github.com/KCCG/mityunder an MIT license.

## 2 Introduction

The human mitochondrial (MT) genome is a 16,569bp circular chromosome, with potentially thousands of copies in a given cell (Davis et al., 2018), and a mutation rate 19x higher than the nuclear genome (Tuppen et al., 2010). Mitochondrial diseases (MDs) are highly heterogeneous genetic disorders, characterised by mitochondrial respiratory chain impairment (Gorman et al., 2016) and are caused by pathogenic variants in either the MT or nuclear genome. Over 300 pathogenic MT variants have been reported (Wallace, 2018), and variants in over 300 nuclear genes associated with disease (Davis et al., 2018). Variants can be present in variable proportions of mitochondrial genomes, known as heteroplasmy. Furthermore, heteroplasmy changes with age and between tissues, being higher in the affected tissue and lower in more accessible tissues such as blood (Sue et al., 1998).

Current clinical-grade whole genome sequencing (WGS) at 30–40× nuclear coverage provides high coverage of the MT genome (1,000–100,000×), suggesting very low levels of heteroplasmy could be reliably detected. With high-coverage however, systematic sequencing errors accumulate, particularly in certain sequence contexts, making it challenging to discern true pathogenic variants from noise (Griffith et al., 2015). Most popular variant callers are optimised for diploid analysis and are thus incapable of identifying low heteroplasmy MT variants. Existing approaches for MT-DNA analysis are web-based (Lee et al., 2008; Weissensteiner et al., 2016) or GUI-based (Ishiya and Ueda, 2017). They are thus less amenable to high-throughput, reproducible analysis, or only validated only on high heteroplasmy variants (Santorsola et al., 2016).

Here, we present *mity*, a bioinformatics pipeline to detect MT SNVs and INDELs from WGS, to assist clinicians and researchers with the diagnosis of MDs. *mity* was optimised to identify low heteroplasmy variants, to be easily integrated into existing high-throughput analysis pipelines, and to generate a highly interpretable report to aid molecular diagnosis.

## 3 Approach

*mity* consists of three modules that easily integrate MT sequence analysis into existing nuclear WGS analysis pipelines (Supp. Fig 1). The first module, *mity-call* analyses a BAM file to call, filter and normalise MT SNVs and INDELs, producing a *mity* VCF. The second module, *mity-report* creates easily interpretable spreadsheet reports with extensive annotations. Finally, the *mity-merge* module merges nuclear and *mity* VCFs to produce a single high-quality VCF. This allows for seamless integration of *mity* into existing production or clinical-grade analysis pipelines for subsequent variant interpretation. *mity* has been optimised for 30–40× Illumina HiSeq X data aligned to the GRCh37 + decoy (hs37d5) reference genome using BWA-MEM.

**Fig. 1.**
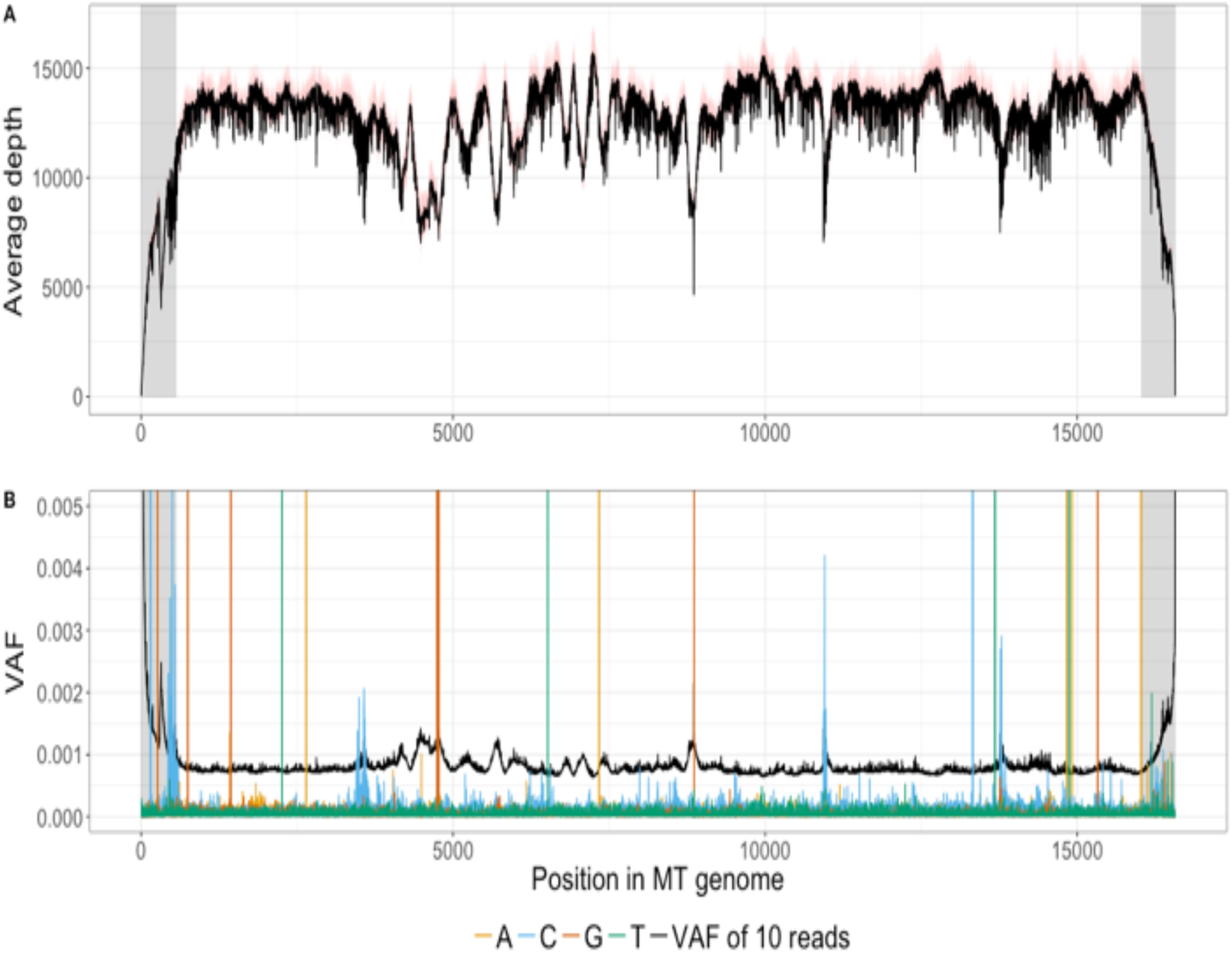
The high average depth across the MT genome in WGS data means that *mity* can detect very low heteroplasmy variants. **A** The average depth (black; red = interquartile range) across 13 replicates of NA12878, with BQ≥24 and MQ≥30. **B** The VAF of all three possible non-reference bases, at each position in the MT genome is typically far lower than the VAF corresponding to 10 high-quality reads (black). Spikes of alternate reads with VAF>0.002 outside the D-loop (grey) correspond to true genetic variants.

### 2.1 mity-call

We first assessed the performance of GATK HaplotypeCaller (Poplin et al., 2018), FreeBayes (Garrison and Marth, 2012), and LoFreq (Wilm et al., 2012) for MT variant detection (Supp. Results). We selected FreeBayes as *mity-call*’s underlying variant caller, as it was more sensitive than HaplotypeCaller, particularly for low heteroplasmy variants and more specific than LoFreq (Supp. Fig 2). Using default FreeBayes settings, the reproducibility from 13 WGS replicates of the NA12878 cell line was poor (Supp. Fig 3a). We thus optimised the mapping quality (MQ≥30), and base quality (BQ≥24) filters, resulting in an average of 17.5±0.5 variants per replicate, with variant allele frequency (VAF) >0.01 (Supp. Fig 3a). These filters caused a minimal reduction of sequencing depth in 2,980 healthy controls (Lacaze et al., 2019; Pinese et al., n.d.) (Supp. Fig 3b). Excluding the D-loop, sequencing depth was generally >10,000× (Figure 1A) in the 13 replicates of NA12878, with low per-base noise, supporting a conservative lower-heteroplasmy limit of >0.002 (Figure 1B). Multi-allelic variants are normalised using a custom algorithm (Supp. Results; Supp. Fig 4).

The default variant quality score in FreeBayes penalises low VAF variants, due to the overwhelmingly high numbers of reference reads. However, we reasoned that the evidence to support low heteroplasmy MT variants should a) only consider the alternate read count, b) scale with the alternate read count, and c) be reported using a similar scale to other variant quality methods. To achieve this, we used a binomial model to implement a Phred-scaled variant quality score, *q*. Assuming a noise level *p, q* is the Phred-scaled probability of observing at least *n* alternate reads by chance, given the total number of reads covering the variant position (Supp. Results). The calculation of *q* is fast, VAF independent, and has a default threshold of *q*≥30, which can be tuned to favour sensitivity or specificity.

*mity* includes three additional filters: a strand bias filter to exclude variants with >90% or <10% alternative reads from one strand, a read depth filter set to <15× depth, and a region filter to exclude variants in the homopolymeric regions at m.302-319, or m.3105-3109, where there is an ‘N’ at m.3107, in the GRCh37 version of the mitochondrial sequence.

### 2.2 mity-report

End-users of *mity* include genome researchers and clinicians, so *mity-report* was developed to produce easily interpretable spreadsheet reports containing comprehensively annotated MT variant lists. Variants are tiered by VAF to aid prioritisation: tier 1, VAF ≥0.01; tier 2, VAF <0.01 with >10 supporting reads; and tier 3 are the remaining variants. Variant impact is estimated using Variant Effect Predictor (VEP) (McLaren et al., 2016), with additional annotation from MITOMAP (Brandon et al., 2005), allele frequency from 2,980 WGS healthy individuals from MGRB and additional VCF fields. The mitochondrial haplogroup is determined using PhyloTree (van Oven and Kayser, 2009).

### 2.3 mity-merge

In order to integrate *mity* into existing WGS analysis pipelines, *mity-merge* replaces the MT variants from a genome-wide VCF (e.g., GATK), with those from the *mity* VCF. The merged VCF can then be used with existing downstream tools for annotation and filtering.

### 2.4 NUMT homology

Nuclear mitochondrial DNA (NUMT) are fragments of the mitochondrial genome integrated into the nuclear genome (Lopez et al., 1994) that can potentially confound heteroplasmy (Parr et al., 2006; Santibanez-Koref et al., 2019). Informed by our previous work in pseudogene homology (Mallawaarachchi et al., 2016), and sequence homology analysis (Supp. Methods & Supp. Results), we conclude that NUMT:MT homology to known NUMT is highly unlikely to cause false-positive tier 1 *mity* variants (Supp. Figure 7). It is challenging to rule out the presence of very rare, or patient-specific NUMT, so we recommend low heteroplasmy variants, particularly tiers 2 and 3 be validated using orthogonal approaches.

### 2.5 Performance

*mity* can operate on either a WGS or MT-only BAM file (<2.1Gb), with a run-time of <10 minutes using a single-core and <8Gb RAM.

## 4 Conclusion

*mity* overcomes many of the challenges of accurate low heteroplasmy variant identification in the MT genome. Additional work is needed to identify MT structural variation, variant phasing, and prioritisation of novel MT variants. *mity* can be easily incorporated into existing high-throughput analysis pipelines, while simultaneously producing user-friendly reports. *mity* was developed on 242 MD patients (Davis et al, in prep) and 2,980 healthy individuals. By extending the scope of variant analysis in patient data, *mity* helps support further adoption of clinical WGS.

## Acknowledgements

We thank members of the Kinghorn Centre for assistance with data generation and patients for contributing their samples to this research.

## 5 Supplementary Methods

### Patient recruitment

We recruited 10 adult patients reviewed at the Mitochondrial Disease Clinic at Royal North Shore Hospital, Sydney, Australia, between 2013-2015. The research was approved by the Northern Sydney Local Health District Human Research Ethics Committee (HREC/10/HAWKE/132). Total genomic DNA was isolated from peripheral blood using standard methods. All participants provided written informed consent. NA12878 reference material was sourced from Genome in a Bottle.

### Sequencing and read alignment

Sequencing libraries were created from nine patients in singlicate, one patient in duplicate, and 13 replicates from NA12878, using Illumina TruSeq Nano HT v2.5 library preparation kits, using Hamilton Star instruments. Sequencing was performed on Illumina HiSeq X instruments following the manufacturers specifications at the Kinghorn Centre for Clinical Genomics, Sydney. Sequence reads were aligned to the human genome reference assembly GRCh37 decoy genome (hs37d5) using BWA-MEM (v0.7.12-r1039, settings -M; (Li, 2013)). Reads were further processed using GATK Indel Realignment, and GATK Base Recalibration (version 3.3; (Van der Auwera et al., 2013)). Depth of coverage was performed using bedtools genomecov (Quinlan, 2014), and depth of alternate alleles in Figure 1B with samtools mpileup (Li et al., 2009).

### Nuclear insertions of mitochondrial origin (NUMTs)

From the comprehensive RHNumtS.2 catalogue of NUMT (Ramos et al., 2011), we identified HSA_NumtS_001 as the only candidate NUMT with sufficient length and homology to be investigated (Supp. Fig 7). We aligned the mitochondrial sequence chrM[GRCh37]:m.5842-9755, corresponding to HSA_NumtS_001 using BLASTN (Altschul et al., 1990), and characterised the distribution of genetic variation within the corresponding NUMT region at chr1[GRCh37]:g.566,391-570,303.

## Supplementary Results

### Evaluation of variant callers

GATK HaplotypeCaller (Poplin et al., 2018) is a popular genome-wide variant caller, but it has three major limitations for MT variant calling: 1) it down-samples the reads to a maximum of 500× depth, 2) it uses a diploid model by default, which is insensitive to low heteroplasmy variants and 3) does not provide a minimum VAF setting. LoFreq (Wilm et al., 2012) was developed specifically for detecting low-frequency variants in somatic cancer data. FreeBayes (Garrison and Marth, 2012) is a popular haplotype-aware, genome-wide variant caller, which allows for control over the minimum VAF and the minimum number of alternative-reads. Whilst FreeBayes and HaplotypeCaller both have ploidy parameters that can theoretically be tuned to prioritise low VAF variants, from a high-ploidy sample, the execution runtime becomes exponentially slower and computationally infeasible in practice for MT analysis.

We assessed the performance of GATK HaplotypeCaller, FreeBayes, and LoFreq for MT variant detection. We first sequenced ten patient genomes using a single lane of Illumina HiSeq X, resulting in 30–40x average nuclear coverage. We ran each of the three callers, on each of the ten samples, and identified a median of 28, 41 and 56 MT variants, respectively (Supp. Fig 1A). We manually inspected every variant and found LoFreq to be overly susceptible to systematic sequencing artefacts, and thus too sensitive. As expected, we found HaplotypeCaller to be insensitive to low VAF variants (Supp. Fig 1B). FreeBayes produced very few false positives and was selected as the *mity-call* variant caller.

### *mity-call*: variant normalisation

As FreeBayes is a haplotype-aware variant caller, it can merge two nearby variants in *cis* into a longer multi-nucleotide variant (MNV). In practice, sequencing noise from high MT coverage can artificially inflate the number of MNVs and reduce the number of reads supporting a variant of interest (Supp. Figure 4). Existing variant decomposition methods including vt normalize (Tan et al., 2015) and vcflib (Garrison, 2016) do not decompose all of the INFO and FORMAT annotations of MNVs, which are critical components for the *mity-report*, and downstream analysis tools. We thus implemented *mity-normalise* to decompose and normalise all variants, as well as the variant metadata within the INFO and FORMAT fields in the VCF (Supp. Figure 4c).

### *mity-call*: assessing variant quality

We implemented a variant quality score, *q*, that is fast and VAF independent, similar to the challenge of identifying rare variants within tumours (Rheinbay et al., 2017). *q* is defined as the Phred-scaled probability of seeing at least the observed number of alternate reads by chance, given a noise threshold and assuming a binomial distribution. That is, given a noise threshold *p* and position *i*, and *n*_*i*_ alternate bases, from a total depth of N_i_, the variant quality q_i_ is:

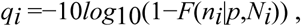

where *F* is the binomial cumulative distribution function. The noise level parameter *p* may vary for each dataset and so should be separately estimated for each dataset.

#### Estimating *p* in the 13 NA12878 replicates

To estimate *p*, we used samtools mpileup (Li et al., 2009) to calculate the VAF of all three alternate bases at every position, in each of the 13 replicates of NA12878 with MQ>30 and BQ>24 (Supp. Figure 5a). There remained a persistent set of alternate reads, with VAF up to 0.001. Thus, taking a conservative approach, we set *p*=0.002 to reduce the impact of sequencing noise. At *q*≥30, we identified 18 variants in all 13 replicates of NA12878 (Supp. Figure 5b). However, there were 5 additional variants seen in a subset of replicates, all of which failed manual review.

#### Estimating *p* in the two replicates of the MD patient

In the same way as above, in two replicates of an MD patient, we estimated a higher noise floor of *p*=0.003 (Supp. Figure 6a) and found 59 variants in both replicates (Supp. Fig 6b), including a known pathogenic m.3243A>G variant (VAF=0.15% in both samples). There were one and three variants private to each replicate, of which two passed manual review, but which just failed the *q* threshold in the other sample.

As with any classification problem, when using *mity* there is a need to balance sensitivity with specificity. We have set the default threshold as *q*≥30, which favours sensitivity over specificity. In larger cohorts, the estimation of *p* could be fine-tuned by estimating a base or region-specific *p*, similar to the ideas presented by Gerstung and colleagues (Gerstung et al., 2014).

### Nuclear insertions of mitochondrial origin (NUMTs)

There have been a number of reports of nuclear mitochondrial DNA (NUMT), which are fragments of the mitochondrial genome integrated into the nuclear genome (Lopez et al., 1994). The presence of NUMT can potentially confound heteroplasmy (Parr et al., 2006; Santibanez-Koref et al., 2019) by causing reads to align to the wrong genome, which would change the variant allele frequency, and thereby the estimation of variant heteroplasmy. Many known NUMT were integrated into the human genome over evolutionary timescales, allowing the accumulation of genetic changes to make it possible to resolve these sequences. To our knowledge, the rates of rare (patient- or family-specific) NUMT have not been systematically investigated and will remain challenging to resolve using short-read sequencing.

Here, we investigated whether any of the known NUMT could create false-positive MT SNV and INDEL variants caused by misalignment of known NUMT reads.

We have previously shown that when using the same sequencing technology (Illumina HiSeq X), and read aligner (BWA-MEM), that 150bp paired-end WGS can correctly align sequencing reads to the *PKD1* gene, despite *PKD1* having six pseudogenes with up-to 97.7% sequence homology (Mallawaarachchi et al., 2016). In a follow-up study of 145 clinical patients, no false positives due to sequencing read misalignment were detected using the same approach (Mallawaarachchi *et al, in review*). These results suggest that any NUMT with homology <97.7% should be readily resolved using 150bp paired-end sequencing reads. We further reasoned that short NUMT less than 300bp, i.e. less than the size of a typical read-pair of 300-450bp, would be easily resolved using existing read alignment. Furthermore, *mity* relies on even more stringent read mapping thresholds than reported above, of MQ>=30, meaning a 1:1000 chance of read misalignment.

Based on the RHNumtS.2 catalogue of NUMT (Ramos et al., 2011), just a single NUMT is longer than 300bp with sequence homology higher than 97.7%: HSA_NumtS_001. This NUMT is 5,839bp long and aligns to chr1:564,464-570,303 (hg19) and chrM:3914-9755 with 98.55% homology. 98.55% homology represents 85 genetic differences over the whole length, or on average, 4.4 mismatches per 300bp read pair.

We performed pairwise sequence alignment of HSA_NumtS_001 and the corresponding MT sequence using BLASTN (Altschul et al., 1990). We reasoned that the genetic differences between the two would allow reads to be properly aligned (Supp Fig 7a), but when the distance between genetic changes approached the read length, that read alignment may become difficult to resolve (Supp Fig 7b). For every base in the alignment, the distance to the nearest mismatch was calculated (Supp Fig 7c) and only one stretch of 24 bases are more than 150 bp from a mismatch, (i.e., the length of a single-read), and none approached 300bp, the paired-end read-length. In this region of high homology, there is still another 150bp read in the read-pair that can be used to align the read pair appropriately

Furthermore, given an average nuclear sequencing depth of 30× and our stringent MQ threshold of 30 (1:1000 read misalignment) even in the unlikely scenario that all of the reads in a given genomic location misalign from the NUMT to the MT, this is highly unlikely to cause a false positive tier 1 variant. Taken together, we find it highly unlikely that given existing 150bp, paired-end reads from WGS, and high-quality read aligners, that NUMT:MT homology to the previously reported NUMT will be a significant source of false positive MT variants.

## 6 Supplementary Figures

**Supp Figure 1:**
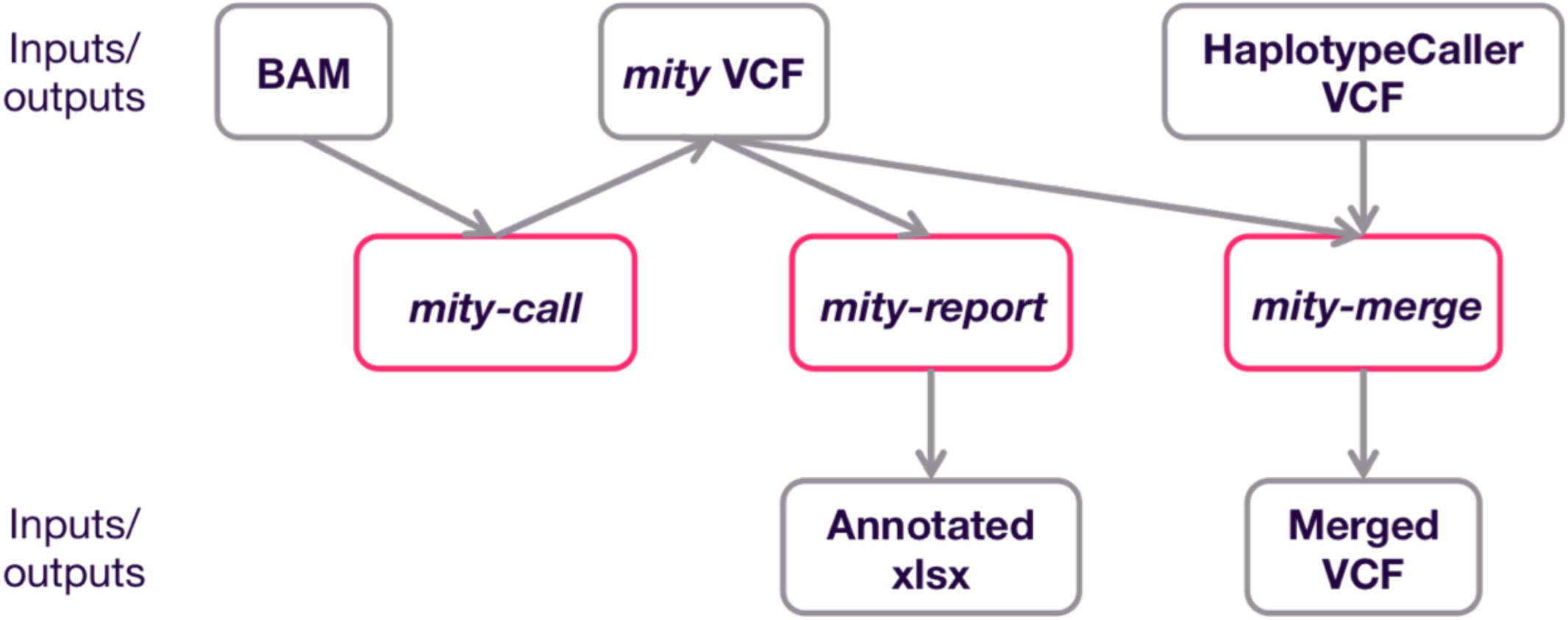
The *mity* analysis pipeline. *mity* consists of three modules: *mity-call*, to call, filter and normalise variants in the MT genome; *mity-report*, to produce a clinician and researcher-friendly annotated MT variant report; *mity-merge*, to integrate *mity* analysis into nuclear analyses.

**Supp Figure 2:**
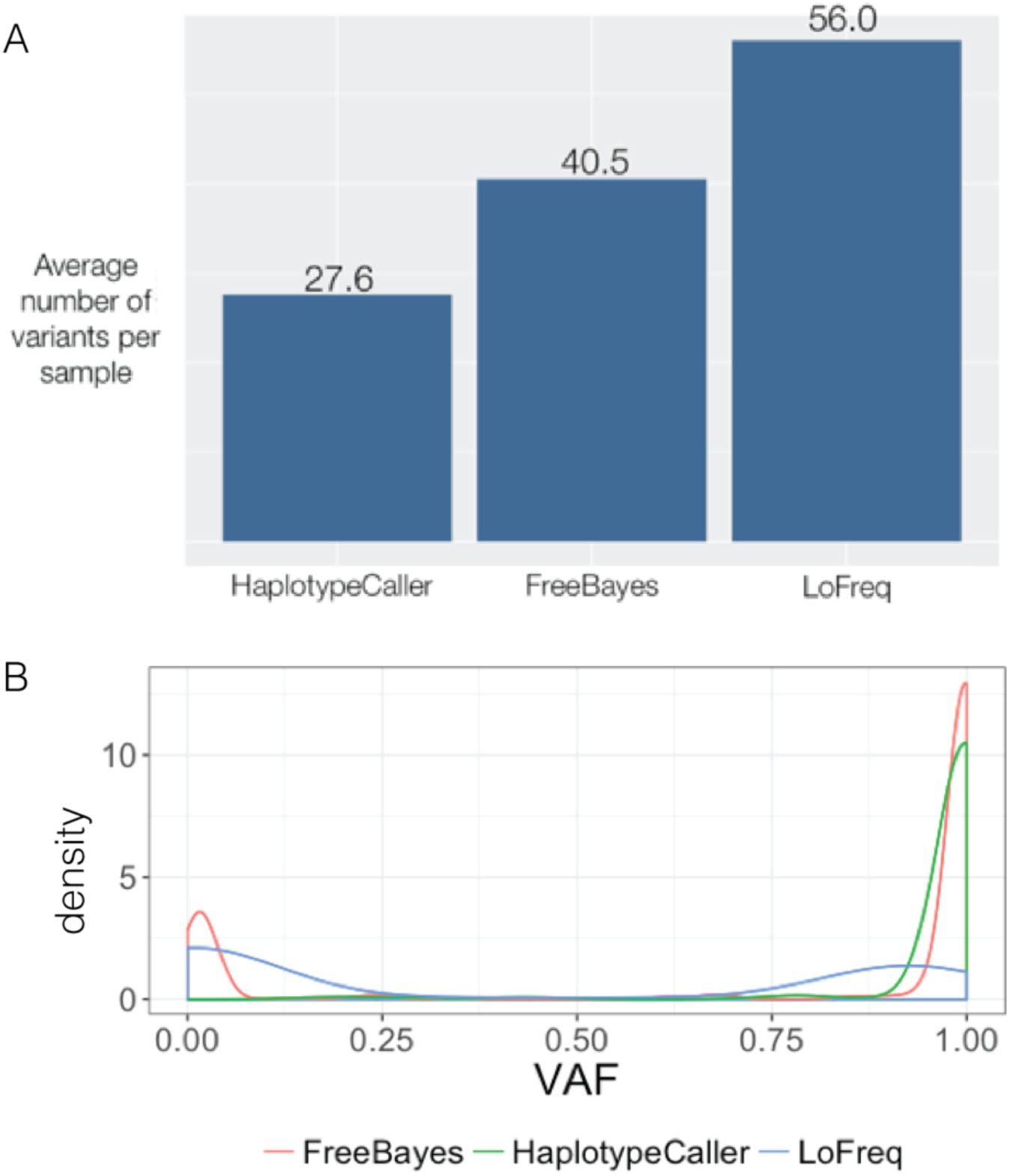
The impact of variant callers on MT variant detection. **A**: Ten patients with MD were sequenced using WGS, and analysed by three different variant callers using default settings: GATK HaplotypeCaller, FreeBayes and LoFreq. The average number of variants found per sample is reported for each caller. **B**: A density plot of the VAF of the variants detected by HaplotypeCaller, FreeBayes and LoFreq. VAF: variant allele frequency.

**Supp Figure 3:**
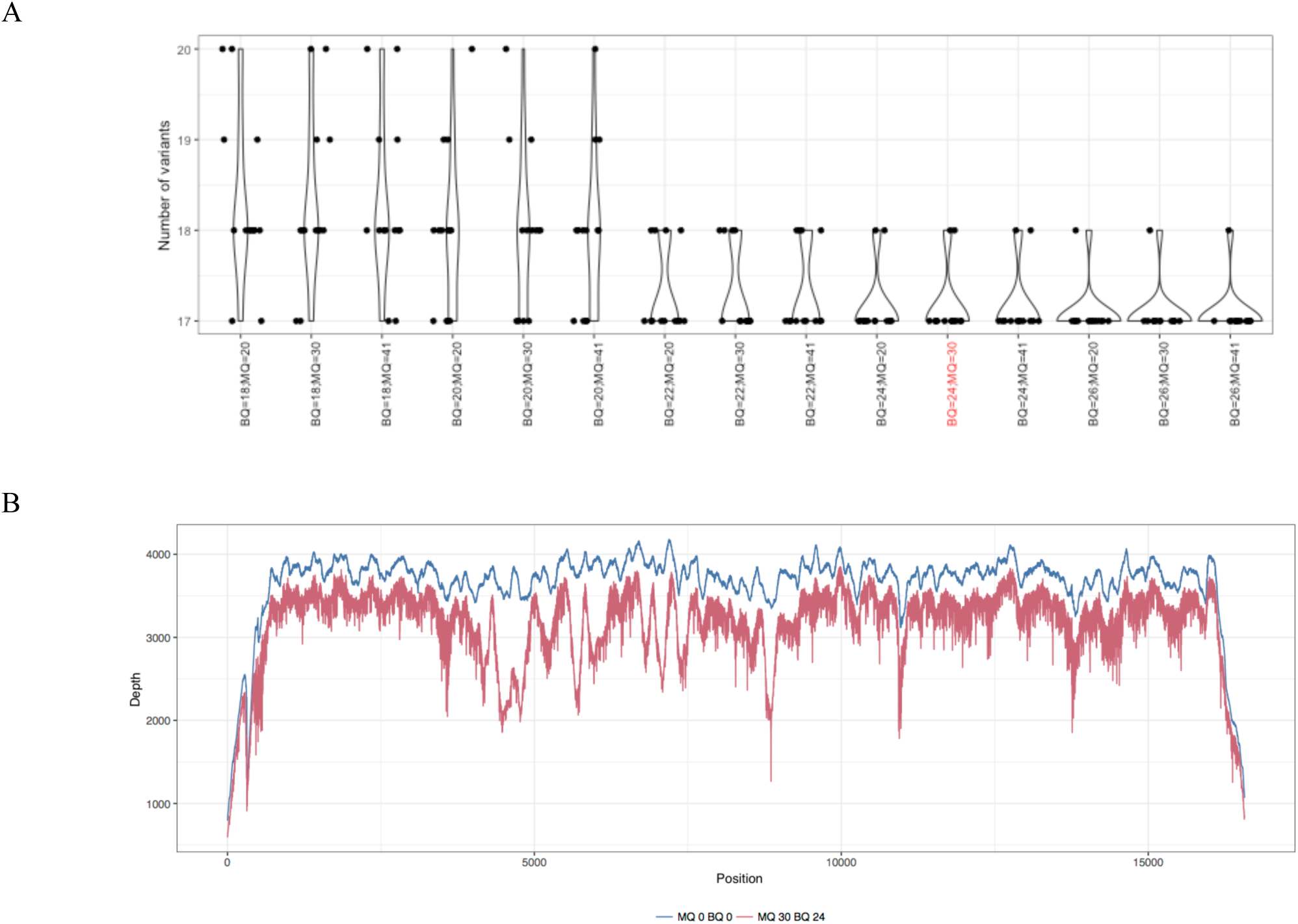
Optimising the mapping (MQ) and base (BQ) quality filters. **A**: The number of variants with VAF>0.01 identified in 13 replicates of NA12878 across varying BQ and MQ combinations. The optimal combination of BQ≤24 and MQ≤30 is highlighted red. **B**: Filtering reads with MQ≤30 and BQ≤24 caused a modest reduction in coverage from 3,666× (MQ=0, BQ=0) to 3,166× (MQ≤30, BQ≤24) in a cohort of 2,980 healthy unrelated controls MGRB. VAF: variant allele frequency; MGRB: medical genomics reference bank.

**Supp Figure 4:**
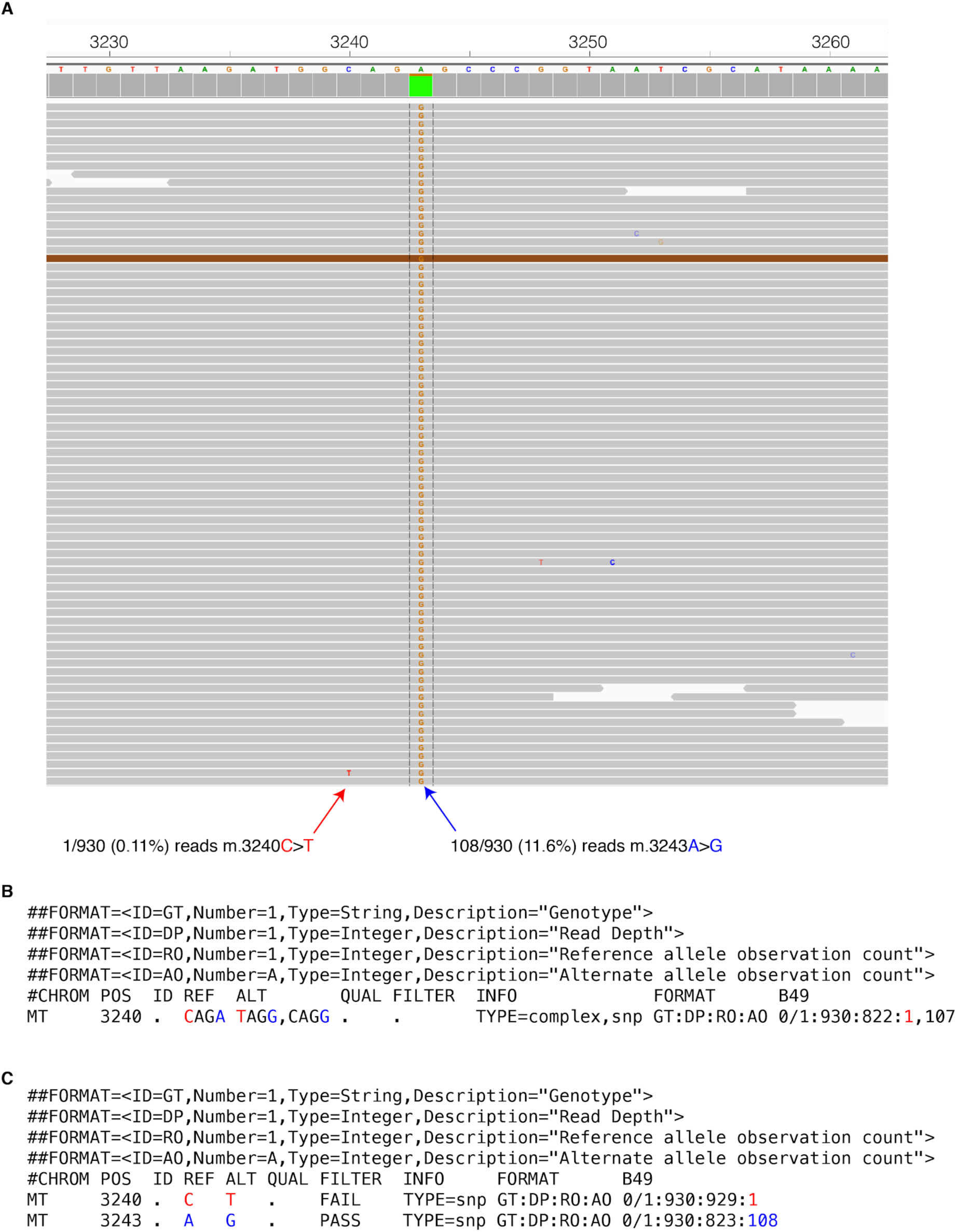
Variant normalisation. **A** Raw sequencing reads from a patient with the pathogenic m.3243A>G variant at 11.6% VAF (blue arrow). A single read had an m.3240C>T variant, with BQ=30 (red arrow). **B** By default, FreeBayes will merge variants on the same haplotype, thus creating a multi-nucleotide polymorphism. Of the 930 total reads, 822 match the CAGA reference sequence, one matches the TAGG sequence, and 107 match the CAGG sequence. Most variant annotation tools, including VEP, which is used by *mity* would fail to annotate this as the well-known pathogenic m.3243A>G variant. **C** After *mity-normalise*, this multi-nucleotide variant is decomposed into the m.3240C>T variant with just one supporting read (red), and the pathogenic m.3243A>G variant with 108 supporting reads (blue). Most variant annotation tools would now easily annotate the m.3243A>G variant as pathogenic. VAF: variant allele frequency, VEP: variant effect predictor, BQ: base quality.

**Supp Figure 5:**
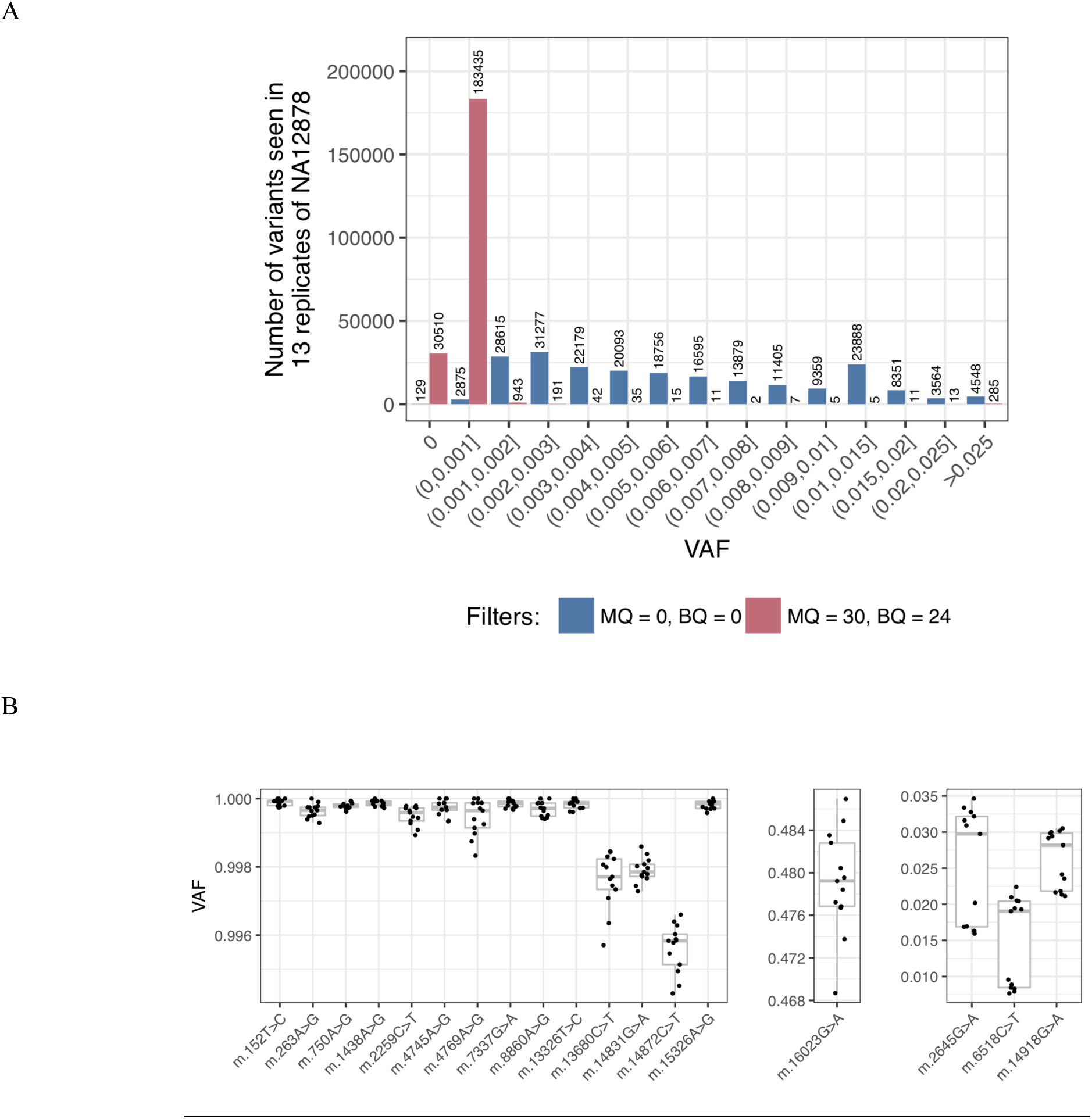
Variant heteroplasmy is highly reproducible using control cell lines. *mity* was run on 13 replicates of NA12878, and we established a noise threshold *p* of 0.002. **A**: The VAF of all alternate alleles found in 13 replicates of NA12878. **B**: The VAF of all *mity* variants identified in all 13 replicates of NA12878, with *p*=0.002, *q*≥30, MQ≥30 and BQ≥24. VAF: variant allele frequency, MQ: mapping quality, BQ: base quality, *p*: noise threshold, *q*: Phred-scaled variant quality.

**Supp Figure 6:**
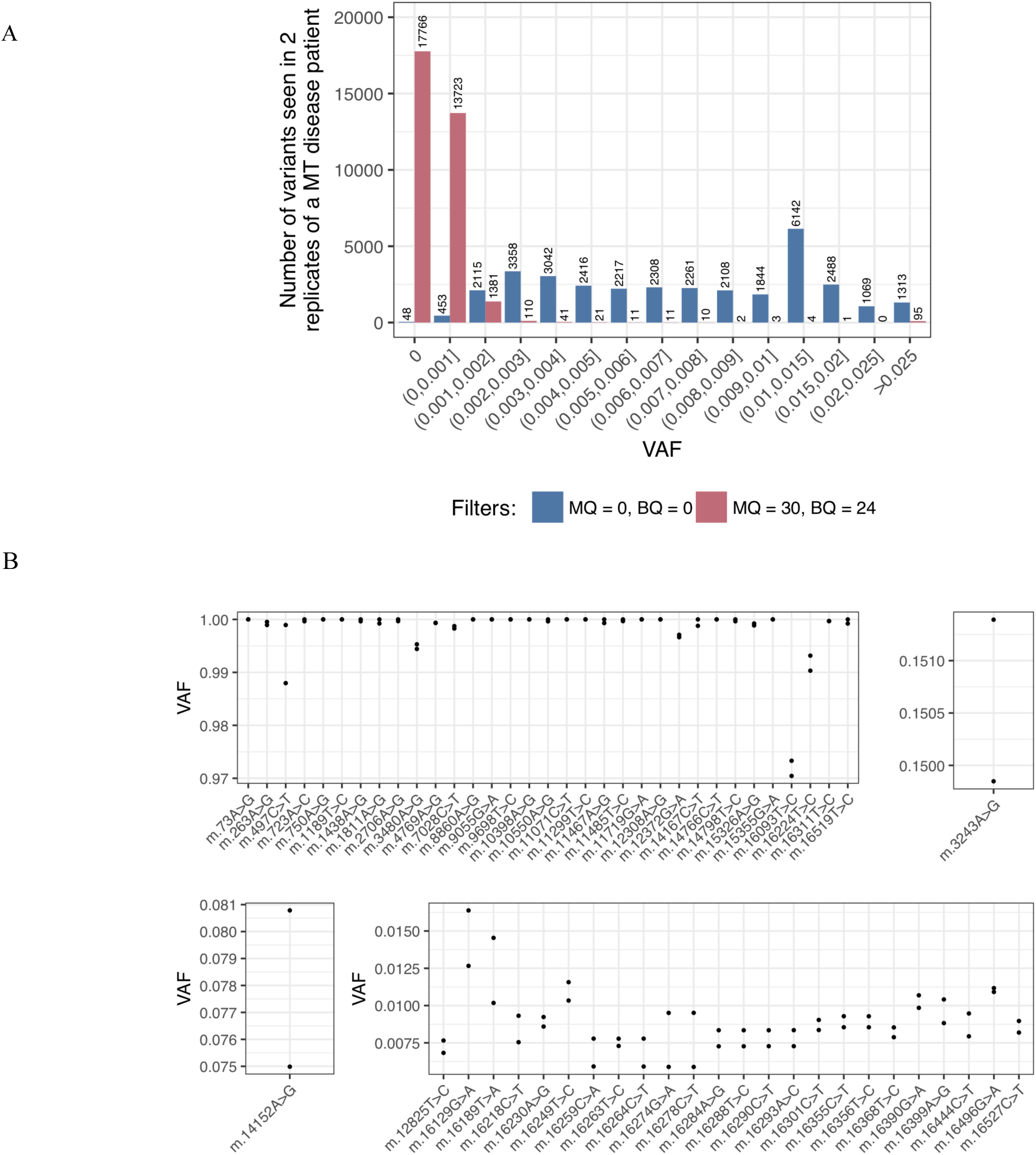
Variant heteroplasmy is highly reproducible using patient material. *mity* was run on two replicates of the same patient with mitochondrial disease. **A**: The VAF of all alternate alleles found in the two replicates. From this, the noise threshold *p* was set to 0.003. **B**: The VAF of all *mity* variants identified in the two replicates, including the m.3243A>G pathogenic variant, with *p*=0.003, *q*≥30, MQ≥30 and BQ≥24. VAF: variant allele frequency, MQ: mapping quality, BQ: base quality, *p*: noise threshold, *q*: Phred-scaled variant quality.

**Supp Figure 7:**
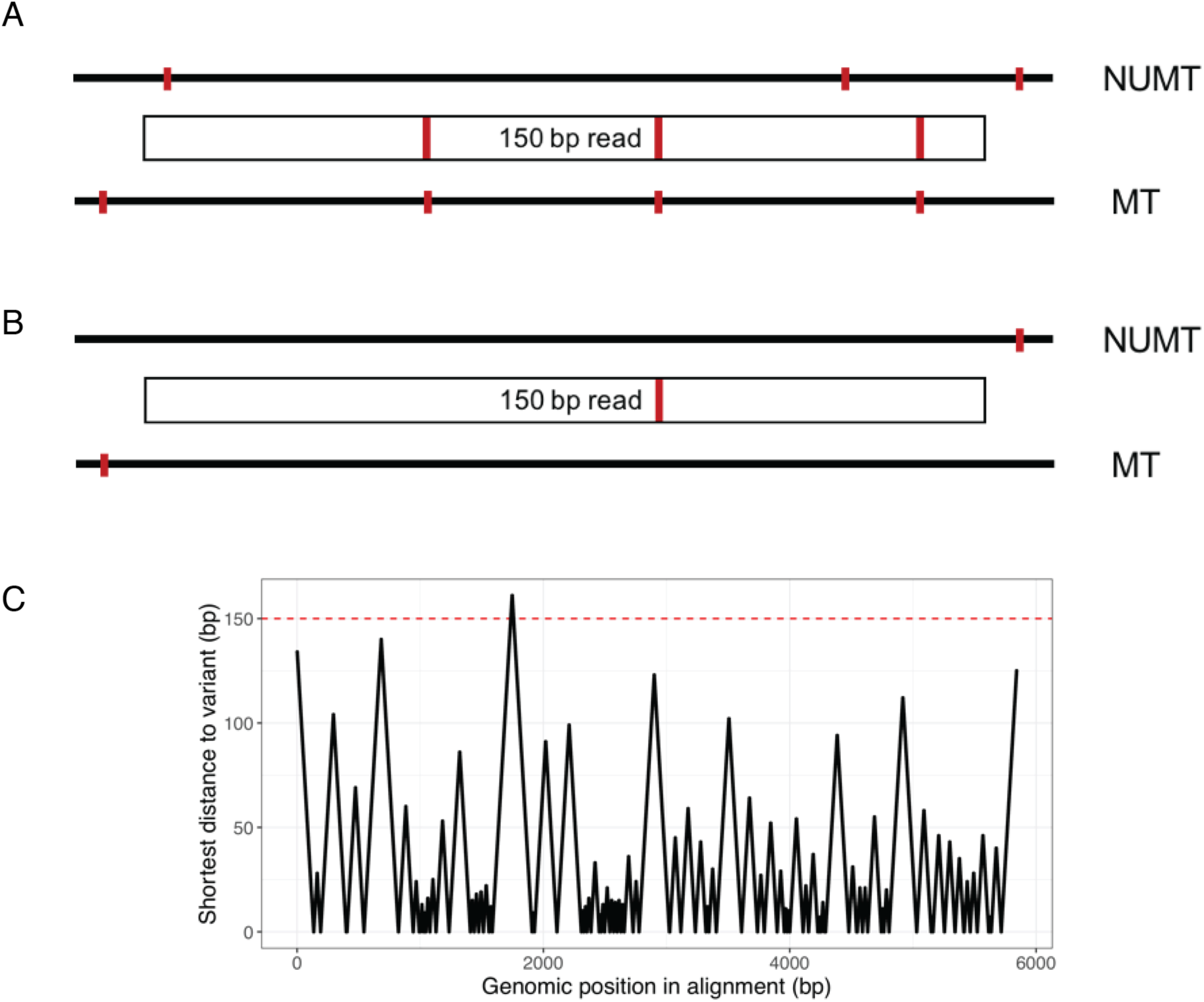
NUMT:MT homology is not expected to be a significant source of false-positive MT variants. Informed by our previous work, we reasoned that reads from any short NUMT (<300bp), or those from a NUMT:MT pair with <97.7% sequence homology, would be correctly aligned and that most NUMT would fall below this homology threshold. In the example shown in panel **A**, the sequence mismatches allow the read to be correctly aligned to the MT. In panel **B**, if the homology is too high, and the read cannot be unambiguously aligned to the MT. **C**: From the RHNumtS.2 catalogue of NUMT(Ramos et al., 2011), there is only one NUMT with length >300bp, and NUMT:MT homology >97%: HSA_NumtS_001. We measured the distance to the nearest mismatch at each position in the alignment of HSA_NumtS_001 and the MT genome using BLASTN. This reveals that there is only a single region of 24 bases where there is more than 150bp between variants. NUMT: nuclear mitochondrial DNA; MT: mitochondrial DNA.

